# Mechanisms of resistance to active state selective tri-complex RAS inhibitors

**DOI:** 10.1101/2025.04.24.649345

**Authors:** Ben Sang, Ling Feng Ye, Feng Hu, Yasin Pourfarjam, Antonio Cuevas-Navarro, Shijie Fan, Zheng Fu, Aaliyah Washington, Diego J. Rodriguez, Alberto Vides, Sumit Kar, Ethan Ahler, Kevin K. Lin, Aparna Hegde, Jacqueline AM Smith, Brian M. Wolpin, Salman R. Punekar, Alexander I. Spira, Ignacio Garrido-Laguna, David S. Hong, Arvin C. Dar, Rona Yaeger, Kathryn C. Arbour, Piro Lito

## Abstract

Tri-complex inhibitors (TCIs) act as molecular glues to recruit cyclophilin A (CYPA) to the active (GTP-bound or ON) conformation of RAS, which in turn prevents the activation of downstream effector proteins like RAF and PI3K. Emerging data demonstrate clinical activity, including tumor regressions, in patients with RAS driven cancers. Despite being promising therapeutic interventions, the mechanisms of resistance in patients treated with these inhibitors remain unknown. Here we studied matched baseline and post-progression specimens from patients treated with the RAS(ON) multi-selective inhibitor daraxonrasib (RMC-6236). Tissue or cell-free DNA specimens were collected from 40 patients with RAS-mutant non-small cell lung, colorectal, or other cancers. Eighteen patients (45%) were found to have acquired alterations in RAS signaling intermediates, including recurrent alterations in KRAS, BRAF, RAF1, MAP2K1/2 and PIK3CA. Preclinical resistance models mirrored the alterations observed in patients. We found that secondary KRAS Y64X mutations caused resistance by disrupting an important pi-pi interaction between KRAS and the indole ring of daraxonrasib, which lowers the affinity of the daraxonrasib:CYPA binary complex for active KRAS. We also identified kinase-dead and low-activity BRAF mutations in samples with acquired resistance. This is puzzling, because TCIs are expected to prevent the interaction between RAS and BRAF, which is needed for hypoactive mutants to dimerize and signal. We now show that RAF dimers are harder to displace from active RAS, as compared to their monomeric forms. Indeed, enhanced RAF dimerization attenuated the ability of TCIs to recruit CYPA to active RAS, resulting in diminished inhibition of oncogenic signaling and tumor growth. Thus, several clinical resistance alterations converge at attenuating the formation of the RAS:daraxonrasib:CYPA tri-complex, either by preventing daraxonrasib binding or by inducing RAF dimers.

## Introduction

The RAS family of GTPases (KRAS, NRAS, HRAS) regulates cell proliferation and survival by cycling between active (GTP-bound, ON) and inactive (GDP-bound, OFF) states^1^. Oncogenic mutations —common in lung, colorectal and pancreatic cancers— impede the ability of RAS to hydrolyze GTP and hyperactivate downstream signaling, leading to uncontrolled growth^2–5^. KRAS G12C selective inhibitors like sotorasib and adagrasib covalently target the inactive conformation of this mutant and are approved by the Food and Drug Administration for the treatment of patients with lung or colorectal cancers (either as monotherapy or combination therapy)^6–12^. While effective in G12C-mutant cancers, the utility of these inhibitors is restricted by their inability to target non-G12C mutants and their inability to target the active form RAS^13,14^.

Tri-complex inhibitors (TCIs) represent a novel class of therapeutics developed in order to overcome these limitations^15–19^. TCIs act as molecular glues to recruit the intracellular chaperone cyclophilin A (CYPA) to the ON state of RAS with varying degrees of mutant selectivity^15,16,20^. The inhibitors elironrasib (RMC-6291) and zoldonrasib (RMC-9805) covalently target KRAS G12C or G12D, respectively, whereas daraxonrasib (RMC-6236) targets multiple RAS variants in a reversible manner^15,16,20^. Regardless of whether RAS engagement is covalent or reversible, the resulting complex of RAS, inhibitor and CYPA (which we refer to as tri-complex) disables oncogenic signaling by preventing typical effector proteins, like RAF and PI3K, from binding to active RAS^15,16^. Daraxonrasib can also stimulate GTP hydrolysis in a RAS mutant selective manner (as recently shown for G12X mutants)^19^. Emerging data in patients with RAS mutant non-small cell lung cancer (NSCLC) and pancreatic cancer treated with RAS(ON) tri-complex inhibitors demonstrate clinical activity, including tumor regressions^21–24^. Clinical activity has also been reported in patients with colorectal cancer (CRC) although these data are less mature^23^. Despite these advances, the mechanisms of resistance in patients treated with RAS(ON) multi-selective inhibitors like daraxonrasib remain under investigation.

## Results

### Alterations associated with daraxonrasib resistance in patients

We evaluated cfDNA and/or tissue specimens from 40 patients with KRAS mutant non-small cell lung (17), colorectal (15), melanoma (4) or other cancers (4) treated with daraxonrasib as part of the Phase 1/1b clinical trial NCT05379985. As shown in Fig. 1A (top heatmap), treatment-emergent alterations in RAS signaling pathway intermediates were identified in 18 patients (45%). Specimens from two patients with KRAS G12D or G12V mutant cancers, were found to have acquired Y64C or Y64D mutations, respectively (Fig. 1A and B). Five patients were found to have treatment-emergent mutations in BRAF. These included kinase-dead (D594A/G/N, Fig. 1A and C) and low-kinase activity (G466E, N581S, K483E) BRAF mutations, which signal in a RAS-dependent manner (and are commonly referred to as Class III BRAF mutations^25–28^), as well as hyper-activating BRAF mutations that signal in a RAS-independent manner, either as monomers (V600E, or Class I) or as dimers (L597V, or Class II)^25–42^. One patient was found to have a mutation in CRAF (or RAF1), in a residue (S257L) that regulates RAF auto-inhibition and dimerization by modulating the interaction with the scaffold protein 14:3:3^43–46^. In addition, SHOC2, which also regulates RAF dimerization^47^, was found mutated in a patient with daraxonrasib resistance. Additional treatment-emergent alterations included KRAS amplification (three patients), MAP2K1/2 mutations (five patients) and alterations in PI3K/MTOR signaling intermediates (four patients). Three patients who did not respond to treatment had baseline alterations in BRAF (E220K, Class III), PIK3CA (E524K) or a CDKN2A deletion (Fig. 1A, bottom heatmap).

**Fig. 1.**
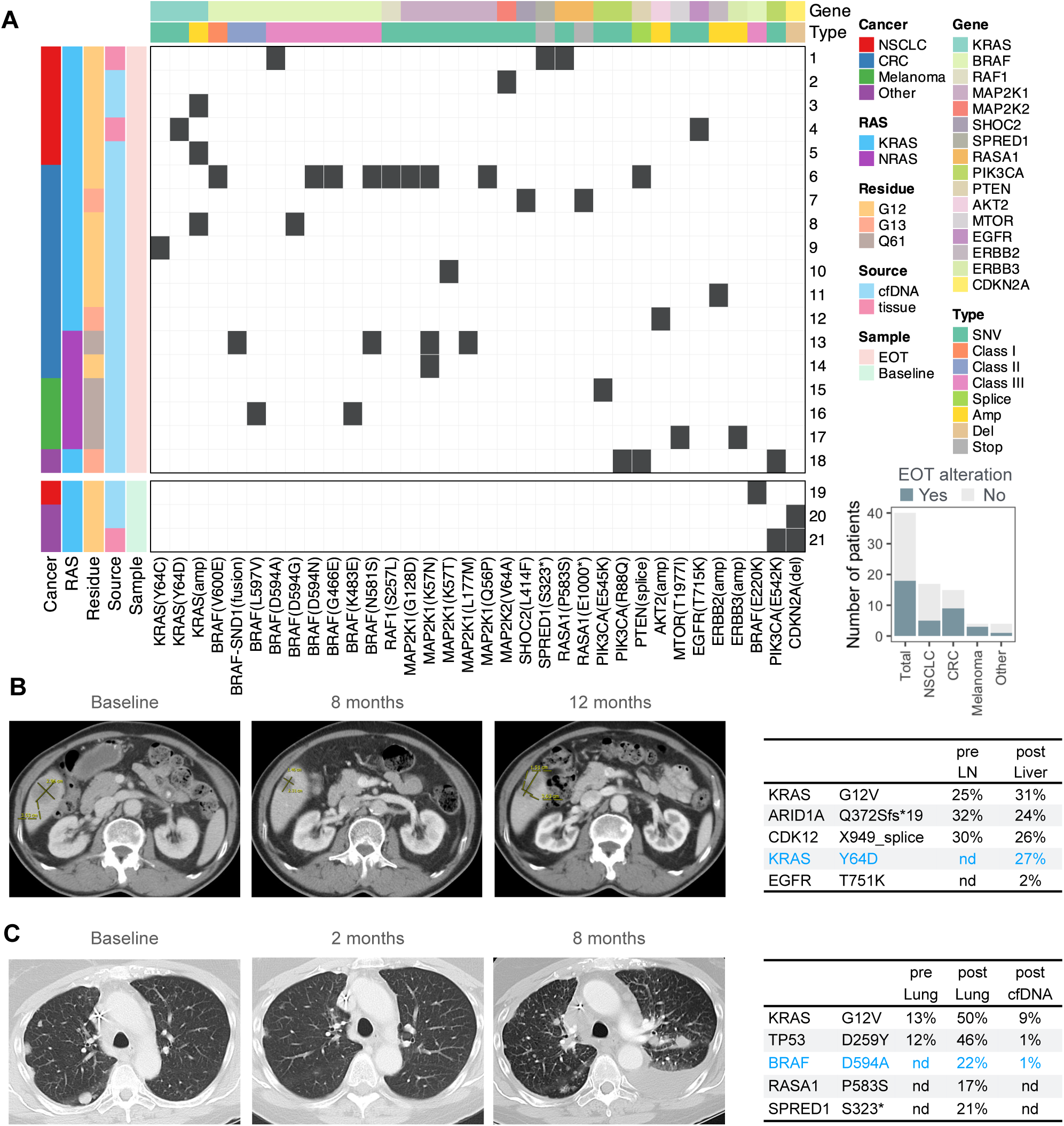
Alterations associated with daraxonrasib resistance in patients. **A**) Heatmap of key treatment-emergent alterations in cell-free DNA (cfDNA) or tissue specimens collected after treatment with daraxonrasib. These alterations were not detected in paired pre-treatment specimens. Bottom heatmap: baseline alterations in select patients who did not respond to treatment. Inset: number of patients with identified alterations in end-of-treatment (EOT) specimens. **B**, **C**) Patients with KRAS^G12V^ mutant lung adenocarcinoma were treated with daraxonrasib and evaluated by computed tomography to determine the effect of treatment over time. The variant allele frequencies of genetic alterations identified in the indicated tissue biopsies at baseline and at daraxonrasib progression are shown. Note the emergence of the secondary KRAS^Y64D^ (**B**) and BRAF^D594A^ (**C**) mutations. LN: lymph node. nd: not detected.

### Resistance alterations in preclinical models

In parallel with the clinical evaluation of resistance described above, we also established preclinical models with acquired resistance, through longitudinal selection of KRAS mutant cell lines with daraxonrasib (or with its related preclinical tool compound RMC-7977). This effort led to the identification of KRAS Y64H and Y71H mutations in KRAS G13D mutant cells (Supplementary Fig. 1A, B) and the identification of KRAS Y64C and BRAF G466A (Class III) mutations in KRAS G12D mutant cells (Supplementary Fig. 1C, D). Temporal passaging of the latter revealed that the Y64C mutation preceded the emergence of the G466A mutation (Supplementary Fig. 1D). Although all resistant passages (R1-5) had a higher daraxonrasib IC_50_, as compared to the parental populations (Par), early passages harboring only the KRAS Y64C mutation (R1 and R2) had a lower IC_50_ than late passages harboring both KRAS Y64C and BRAF G466A (R4 and R5). The variant allele frequencies of the KRAS G12D (∼100%) and Y64C (∼50%) mutations in Supplementary Fig. 1D, suggest the presence of a doubly mutated KRAS protein.

### Y64 mutations drive resistance by attenuating tri-complex formation

We next sought to independently validate if the treatment-emergent alterations drive resistance and to determine the mechanism by which they do so. We first hypothesized that Y64 mutations (Fig. 1b) interfere with the formation of the KRAS:daraxonrasib:CYPA tri-complex, which is necessary for blocking effector signaling downstream of mutant KRAS. Indeed, the crystal structure of KRAS G12D in complex with the RAS(ON) multi-selective inhibitor RMC-7977 revealed that Y64 forms a pi-pi stacking interaction with the indole ring of the TCI (Fig. 2A). To experimentally determine the effect of Y64 mutants on daraxonrasib, we used HEK293T cells expressing KRAS and CYPA or the RAS-binding domain (RBD) of CRAF (CRAF^RBD^), as part of a split luciferase complementation reporter (see schematics in Supplementary Fig. 1E and F). Binding of KRAS to CYPA or to CRAF^RBD^ reconstitutes the luciferase enzyme, permitting the quantification of these complexes in live cells. As expected, Y64D/C/H mutations attenuated the ability of daraxonrasib to stimulate the formation of the KRAS-CYPA complex (Fig. 2b and Supplementary Fig. 1E). In turn, this led to attenuated displacement of the CRAF^RBD^ from oncogenic KRAS (Fig. 2C and Supplementary Fig. 1F), coupled with attenuated inhibition of downstream signaling (Fig. 2D and Supplementary Fig. 1G). Of note, daraxonrasib was able to partially stimulate CYPA binding to G12D/Y64X mutant KRAS.

**Fig 2.**
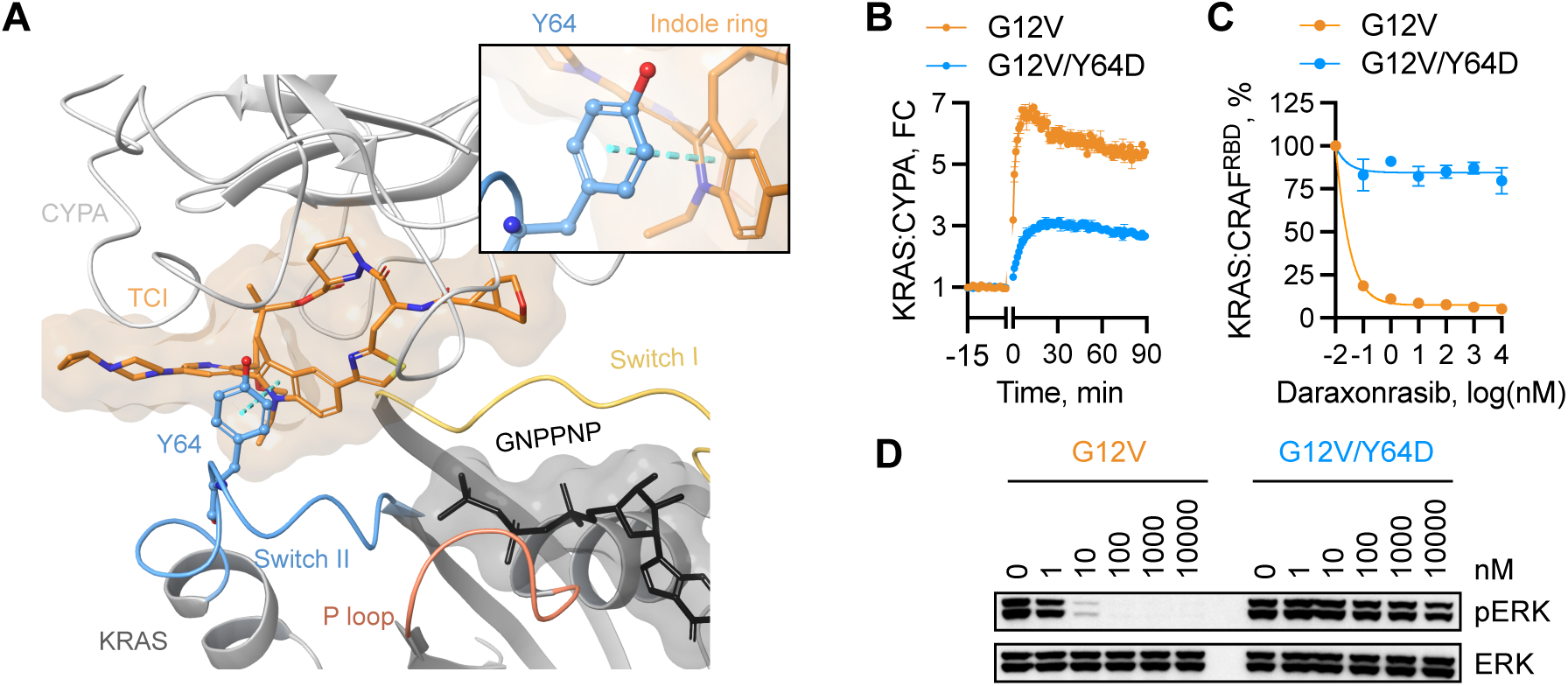
Y64 mutations attenuate the recruitment of CYPA to active KRAS. **A**) Crystal structure of KRAS^G12D^ in complex with RMC-7977 and CYPA. The dashed line indicating a pi-pi interaction between KRAS Y64 and the indole ring of RMC-7977. **B**) Live cells expressing a split-luciferase reporter detecting the complex between CYPA and the indicated KRAS mutants (see schematic in Supplementary Fig. 1E) were treated with 100 nM daraxonrasib and assayed over time, before and after treatment. FC: fold change. **C**) As in **B**, but with a reporter detecting the complex between the CRAF RBD and mutant KRAS (see schematic in Supplementary Fig. 1F). **D**) Extracts from HEK293T cells expressing the indicated single- or Y64 double-mutants treated with daraxonrasib (2 h) were analyzed to determine the effect on ERK activation. In **B** and **C**, n = 3, mean ± s.e.m. A representative of two independent experiments is shown in **D**.

### Kinase-dead or impairing BRAF mutations drive resistance

The identification of the kinase-dead (D594A, Fig. 1C) and other hypoactive Class III BRAF mutations (Fig. 1A) in patients whose tumors progressed on daraxonrasib is puzzling. These mutants function as obligate dimers^25,26,30,48–50^, i.e., they require RAS-dependent dimerization with wild-type RAF kinases, in order to activate downstream ERK signaling. RAS(ON) multi-selective TCIs like daraxonrasib, however, are expected to block RAS-induced BRAF dimerization. Thus, it is not clear if (and how) Class III BRAF mutations drive daraxonrasib resistance, or if these are instead passenger mutations.

To test this, wild-type or kinase-dead BRAF were expressed in KRAS G12V mutant lung (H727) or colorectal (SW620) cancer cells under a doxycycline (dox)-induced promoter. In both models, kinase-dead, but not wild-type BRAF, led to an attenuation of the effect of daraxonrasib on ERK signaling (Fig. 3A and Supplementary Fig. 2A, B) and proliferation (Fig. 3B and Supplementary Fig. 2C-F). An attenuated inhibition was also observed for other Class III BRAF mutants (Supplementary Fig. 3). The attenuated inhibition caused by kinase-dead BRAF in lung and colorectal cancer models was reversed when secondary mutations that disrupt dimerization (i.e., R509H) or RAS-binding (i.e., R188L) were introduced alongside D594A (Fig. 3A, B and Supplementary Fig. 4A, B). Expression of kinase dead BRAF also attenuated the effect of covalent G12C(ON) tri-complex inhibitors (Supplementary Fig. 4C and D).

**Fig 3.**
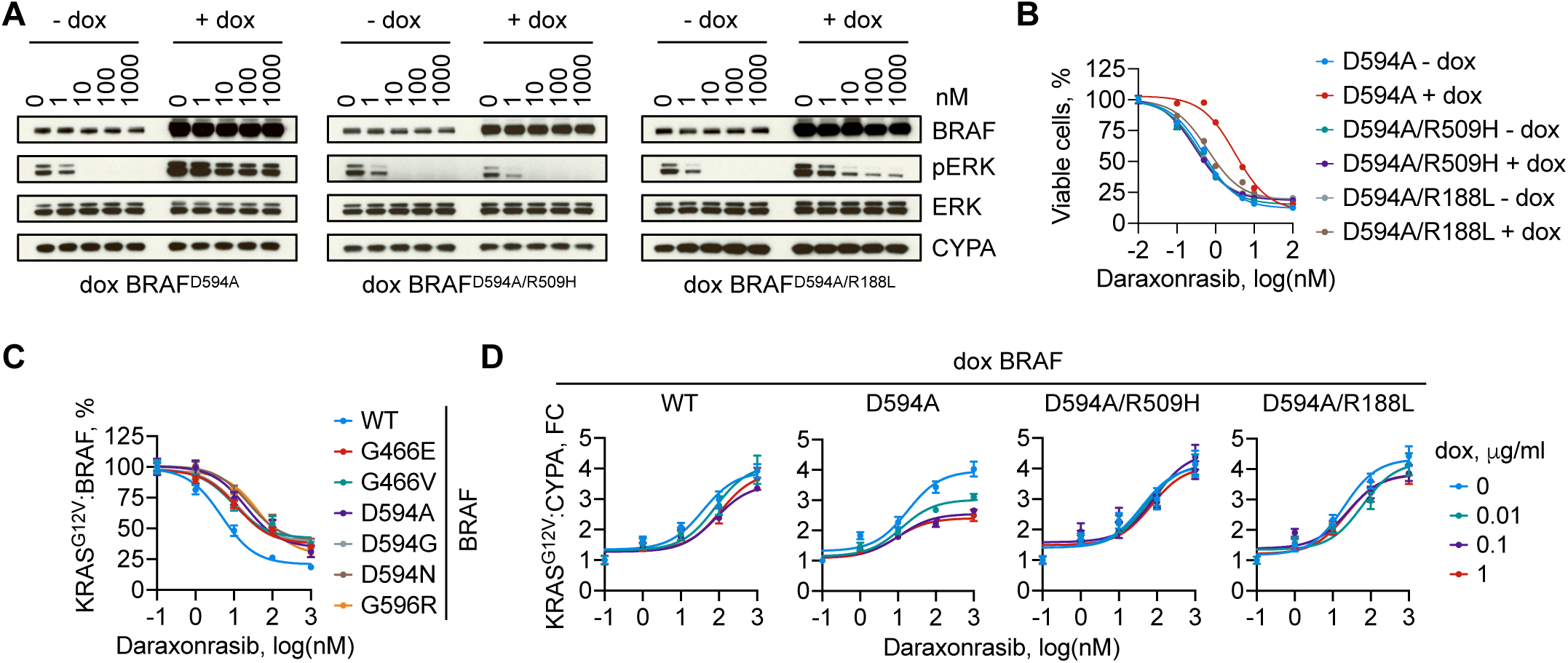
Kinase-dead or impairing BRAF mutations drive daraxonrasib resistance by attenuating target engagement. **A**, **B**) KRAS^G12V^ mutant (H727) lung cancer cells expressing the indicated BRAF mutants under a dox-inducible promoter were treated with increasing concentrations of daraxonrasib for 2 h to determine the effect on ERK signaling (**A**) or for 72 h to determine the effect on cell viability (**B**). **C)** Live cell split-luciferase assay detecting the effect of daraxonrasib on the interaction between KRAS^G12V^ and the indicated BRAF variants. **D**) Live cell split-luciferase assay detecting the effect of dox-inducible expression of the indicated BRAF mutants on the ability of daraxonrasib to recruit CYPA onto KRAS^G12V^. FC: fold change. The daraxonrasib treatment time in **C** and **D** is 2 h. A representative of two independent experiments is shown in **A**. In **B**-**D**, n = 3, mean ± s.e.m.

Daraxonrasib was less effective at displacing kinase-dead and other hypoactive Class III BRAF mutants from oncogenic KRAS, as compared with wild-type BRAF (Fig. 3C and Supplementary Fig. 4E). Moreover, the ability of daraxonrasib to recruit CYPA to oncogenic KRAS was attenuated with increasing amounts of kinase-dead (but not wild-type) BRAF, an effect that was again dependent on intact dimerization (Fig. 3D and Supplementary Fig. 4C: note the KRAS electrophoretic motility shift). Together, these data suggest that hypoactive (Class III) BRAF mutants drive daraxonrasib resistance by impeding target engagement and inhibition.

### Attenuated displacement of RAF dimers from mutant KRAS by daraxonrasib

The data above suggest that dimerization is the underlying mechanism that enables hypoactive BRAF mutants to antagonize the effect of daraxonrasib. To directly test this possibility, we used a chemical-ligand induced dimerization system^51^ that uncouples dimerization from RAF mutations (see schematic in Supplementary Fig. 5A). In this system, RAF proteins are tagged with DmrA or DmrC, which can then be induced to dimerize via an exogenous ligand (A/C ligand). A/C ligand-induced BRAF:CRAF, BRAF:BRAF and CRAF:CRAF dimers had an attenuated displacement from oncogenic KRAS by daraxonrasib (Supplementary Fig. 5B, D, F: compare A/C ligand vs DMSO). In agreement, ligand-induced dimerization led to diminished KRAS:CYPA complex formation by daraxonrasib (Fig. 4A), coupled with attenuated ERK signaling inhibition (Supplementary Fig. 5C, E, G) and antiproliferative effect in KRAS G12V mutant lung and colorectal cancer cells (Supplementary Fig. 5H). A similar effect was observed across various KRAS mutants and inhibitors (Supplementary Fig. 6). Thus, RAF dimerization phenocopies the effect of several mutations found in patients treated with daraxonrasib.

**Fig 4.**
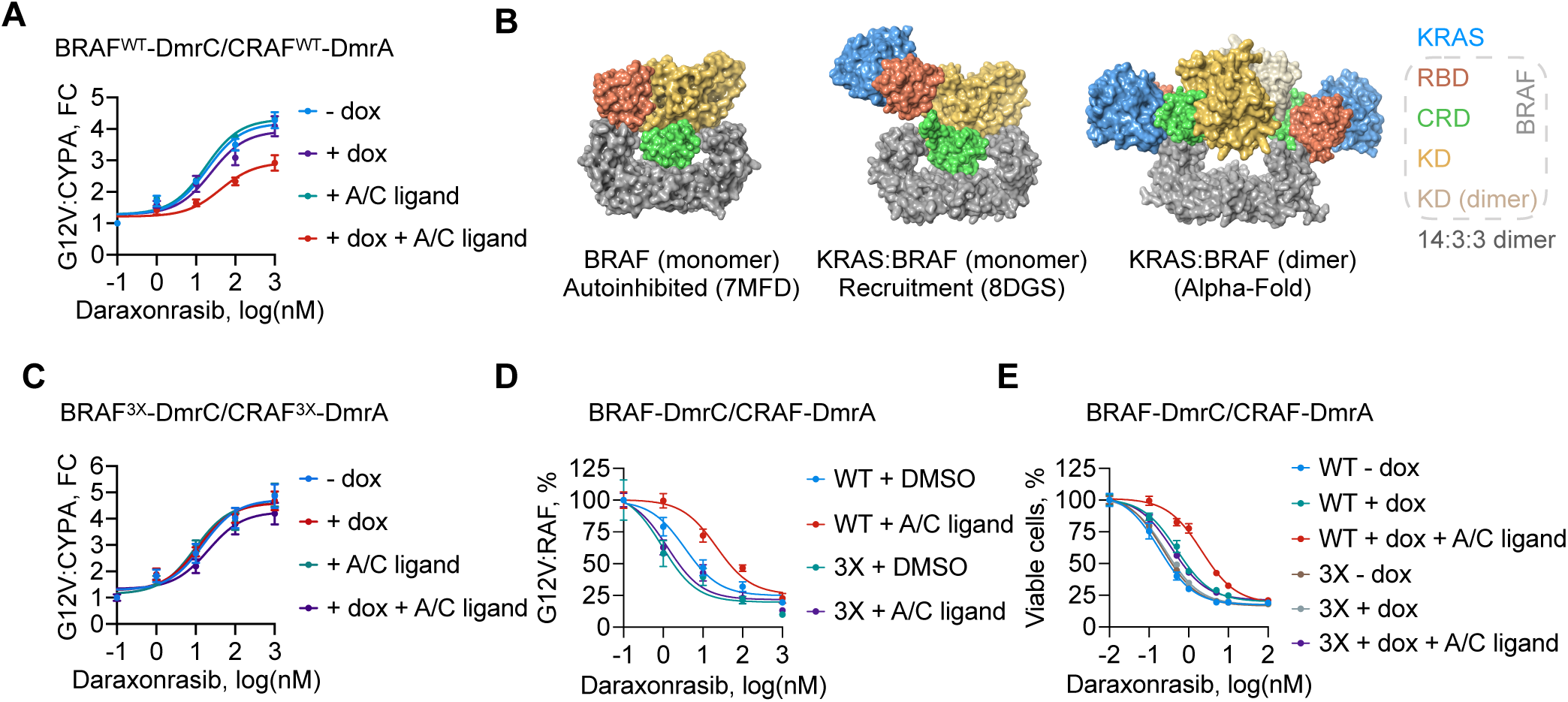
RAF dimerization attenuates target engagement via the cysteine-rich domain. **A**) Live cell split-luciferase assay detecting the effect of dox-inducible expression of the indicated BRAF/CRAF ligand-induced dimerization constructs on the ability of daraxonrasib (2 h) to recruit CYPA onto mutant KRAS. The A/C ligand induces BRAF:CRAF dimerization via the DmrA/C domains. **B**) Models of BRAF in complex with the scaffold protein 14:3:3 in auto-inhibited state (left) or in complex with 14:3:3 and KRAS, either as a monomer (middle) or dimer (right). **C**) As in **A** but with BRAF/CRAF ligand-induced dimerization constructs harboring CRD mutations (3X) that impede KRAS binding. BRAF^3X^: L232A/ F237A/E275A, CRAF^3X^: L136A/F141A/K179A. **D**) Live cell split-luciferase assay detecting the effect of A/C ligand induced dimerization of the indicated BRAF/CRAF constructs on the ability of daraxonrasib to inhibit the interaction between KRAS^G12V^ and RAF. See also Supplementary Fig. 7C. **E**) KRAS^G12V^ mutant (H727) lung cancer cells expressing the indicated constructs under a dox-inducible promoter were treated as shown for 72 h to determine the effect of the indicated treatments and CRD mutations on cell viability. FC: fold change. In **A**, **C**, **D**, **E**, n = 3, mean ± s.e.m.

Although the structural basis of how RAS interacts with the RBD and the cysteine-rich domain (CRD) of RAF is known^52^, there are no high-resolution structures to inform how this interaction is coordinated when RAF is dimerized. It has been suggested that for dimerization to occur, the CRD of RAF must be displaced from its auto-inhibited conformation^53–57^ (where it is found sandwiched between the kinase domain of BRAF and 14:3:3, Fig. 4B: left and middle). To better understand how RAF dimers might affect daraxonrasib’s ability to recruit CYPA to active RAS, we used Alpha-Fold^58^ (AF) to model the KRAS:BRAF:14:3:3 homodimer complex and found that both RBD and CRD were complexed with KRAS. The latter prediction was similar to the configuration of the KRAS-CRAF^RBD-CRD^ interaction, as evidenced in recent crystal structures^52^.

The observations above suggest that enhanced RAF dimerization attenuates the ability of daraxonrasib to recruit CYPA to mutant KRAS in a CRD-dependent manner. Indeed, daraxonrasib could displace the RAF^RBD^ from oncogenic KRAS with a lower IC_50_, as compared to the RAF^RBD-CRD^ (Supplementary Fig. 7A), an observation that was true for either BRAF or CRAF. Mutations in three CRD residues that are necessary for maximal KRAS binding (hereafter referred to as 3X), enhanced the ability of daraxonrasib to displace RBD-CRD from mutant KRAS (Supplementary Fig. 7B). More importantly, these mutations could reverse the effect of ligand-induced RAF dimerization on the ability of daraxonrasib to recruit CYPA onto mutant KRAS (Fig. 4c) and displace RAF dimers from it (Fig. 4D), as well as on the effect of daraxonrasib on the inhibition of ERK signaling (Supplementary Fig. 7C and D) and proliferation (Fig. 4E and Supplementary Fig. 7E).

## Discussion

Our study thus identifies mechanisms of resistance in patients with KRAS mutant cancers treated with daraxonrasib, a RAS(ON) multi-selective tri-complex inhibitor. Our findings highlight multiple resistance pathways, including secondary RAS mutations, alterations in RAF kinase function, and other MAPK and/or PI3K pathway alterations. One resistance mechanism involves secondary mutations in KRAS, specifically at Y64, which directly interfere with tri-complex formation. Structural and biochemical analyses demonstrate that Y64 engages in pi-pi stacking interactions with the indole ring of daraxonrasib, an important determinant for stable tri-complex formation. This alteration was observed both in patients and in pre-clinical models. In the latter setting, Y64 mutations emerged in a stepwise manner alongside other alterations, suggesting a coordinated evolutionary process driving resistance.

A second major resistance mechanism involves alterations in RAF kinases. Class I and Class II BRAF mutants have high kinase activity and signal independently of RAS. These are highly likely to drive resistance by bypassing oncogenic KRAS and were not studied in detail here. The identification of kinase-dead and other low-activity Class III BRAF mutations in patients whose tumors progressed on daraxonrasib is surprising. These mutants rely on RAS-dependent dimerization with wild-type RAF proteins to sustain ERK signaling. However, because RAS(ON) inhibitors are designed to block RAS-RAF interaction, which is a necessary step for RAF dimerization, the persistence of ERK signaling in this setting is unexpected. Our functional studies reveal that hypoactive BRAF mutants attenuate the effect of daraxonrasib by impairing target engagement in a dimerization-dependent manner. Consistent with this, ligand-induced RAF dimerization phenocopied the effect of these mutations, attenuating the KRAS-daraxonrasib-CYPA tri-complex formation and reducing daraxonrasib-mediated ERK pathway inhibition. Structural modeling suggests that RAF dimerization repositions the CRD in a manner that enhances KRAS binding, making it more resistant to displacement by tri-complex inhibitors. Indeed, mutating CRD residues that are key for optimal KRAS binding restored daraxonrasib sensitivity in dimerized RAF contexts, providing further evidence that enhanced RAF dimerization directly impedes the ability of various inhibitors to inactivate mutant RAS.

These findings have significant therapeutic implications. First, the emergence of Y64 mutations suggests that next-generation tri-complex inhibitors may need to accommodate or counteract this structural alteration to maintain efficacy. Second, the ability of hypoactive RAF mutants to drive resistance suggests that other acquired genetic alterations (e.g., CRAF, ARAF and SHOC2) or adaptive signaling changes (e.g., loss of ERK-dependent feedback), which trigger RAS-dependent dimerization can antagonize the effect of daraxonrasib treatment. Together, these possibilities underscore the need for combination strategies that can effectively block RAF dimerization or exploit vulnerabilities in dimer-dependent signaling. One potential approach could involve combining daraxonrasib with dimer-disrupting RAF inhibitors^59^ or direct ERK inhibitors to overcome resistance in tumors where ERK signaling is driven by RAF dimerization. Third, our insights suggest that disrupting CRD-KRAS interactions could be a viable strategy to enhance the displacement of RAF dimers by RAS(ON) inhibitors. Overall, our study provides a mechanistic framework for understanding acquired resistance to daraxonrasib in KRAS-mutant cancers.

## Methods

No statistical methods were used to predetermine sample size. The experiments were not randomized and, unless otherwise indicated, the investigators were not blinded to allocation during experiments and outcome assessment.

### Patient specimens

Patients were eligible if they were treated with daraxonrasib (clinical trial: NCT05379985). Most pre-treatment and post-treatment specimens were obtained through liquid biopsies. Tissue biopsies (2 patients) or cfDNA (rest of patients) were analyzed for genetic alterations. The GuardantINFINITY assay (Guardant Health, Palo Alto, CA) was used to for next-generation sequencing (NGS) of research specimens collected from the above study. Additionally, targeted NGS of clinical specimens was carried out independently of this study as part of routine clinical assessment with established point-of-care assays: MSK-ACCESS for liquid biopsies, and MSK-IMPACT for tissue biopsies. Alterations in genes outside these targeted panels could not be assessed. The data were collected through chart review or clinical trial repository. Treatment-emergent variants were defined as those detected only after exposure to therapy. The daraxonrasib clinical trial (NCT05379985) is being conducted in accordance with recognized U.S. ethical guidelines (i.e., U.S. Common Rule) and per local institutional review board guidelines. All patients included in the clinical trial were subject to and provided written informed consent prior to study enrollment. Daraxonrasib was administered once daily in 21-day cycles to patients enrolled on the protocol.

### Targeted bulk exome sequencing

Genomic DNA (gDNA) extracted from cell line models was subjected to targeted-capture massively parallel sequencing using the Memorial Sloan Kettering–Integrated Mutation Profiling of Actionable Cancer Targets (MSK-IMPACT) sequencing assay, as previously described^60,61^. In brief, all protein-coding exons and selected introns of 550 commonly implicated oncogenes, tumor suppressor genes and genes encoding members of pathways deemed actionable by targeted therapies were captured by hybridizing barcoded libraries to designed custom oligonucleotides (Nimblegen SeqCap). gDNA (100–250 ng) was used to prepare barcoded sequence libraries (Kapa Biosystems), which were combined into equimolar pools of 13–21 samples. These pools were subsequently subjected to Illumina HiSeq 2000 for paired-end 100-bp sequencing, which produced a median of 588-fold coverage per DNA sample. CASAVA was used to demultiplex sequenced data, and reads were aligned to the reference human genome (hg19) using BWA and post-processed using GATK according to GATK best practices. Single-nucleotide variants and small insertion–deletions (indels) were called through MuTect and GATK, respectively. The sequencing and data analysis was carried out in our institutional core facility. DNA from a pool of unmatched normal samples was used as normal baseline for comparison with sequenced samples to eliminate common polymorphisms and systematic sequencing artifacts.

### Cell culture and reagents

The NCI-H727 (CRL-5815), SW620 (CCL-227), NCI-H358 (CRL-5807), AsPC-1 (CRL-1682), LoVo (CCL-229) and 293T (CRL-11268) cell lines were obtained from American Type Culture Collection (ATCC), confirmed by STR profiling, and tested negative for mycoplasma. All cell lines were maintained in DMEM medium supplemented with 10% FBS, 2 mM L-glutamine, penicillin, and streptomycin. The RAS inhibitors daraxonrasib, RMC-7977, RMC-4998 and RMC-9945 were provided by Revolution Medicines.

### Resistant models from cell culture

Cells were cultured in the presence of increasing concentrations (10–300 nM) of daraxonrasib (or RMC-7977) and passaged when reaching approximately 80% confluence. The selection process spanned 4-20 months. Following selection, resistant cell lines were maintained in daraxonrasib. Serial sampling of drug-selected populations was carried out at various passages to determine the sensitivity to daraxonrasib as well as emerging genetic alterations, by using the assays described below. Serial passages were further expanded (under the original selection condition) for downstream experiments. The passage number indicated in the figures does not reflect the same time interval between passages or lineages.

### Immunoblotting and antibodies

Cells were collected and lysed with lysis buffer (50 mM Tris-HCl pH 7.5, 1% NP-40, 150 mM NaCl, 10% glycerol and 1 mM EDTA) supplemented with Halt protease and phosphatase inhibitor cocktail (Thermo Fisher Scientific) on ice for 10 min. Lysates were then centrifuged at 16,000g for 10 min, followed by protein quantification using the Pierce BCA Protein Assay kit (Thermo Fisher Scientific). Protein samples were prepared in 1X SDS loading buffer and heated at 95 °C for 5 min. 30 μg of total protein was loaded to each well of 4–12% SDS–PAGE gels (Thermo Fisher Scientific). The gels were run in 1X MOPS buffer (Thermo Fisher Scientific) at 100 V for 2 h. Proteins were then transferred to nitrocellulose membranes (GE Healthcare) with 1X Tris-glycine buffer (Bio-Rad) at 100 V for 1 h. After blocking with 5% skim milk for 1 h at room temperature, membranes were incubated with primary antibodies overnight at 4 °C. Detection was performed using HRP conjugated secondary antibodies and ECL substrate (Thermo Fisher Scientific).

Primary antibodies used include: BRAF (1:2000, sc-9002, Santa Cruz Biotechnology), CYPA (1:1000, sc-134310, Santa Cruz Biotechnology), Phospho-ERK (1:1000, 9101, Cell Signaling Technology), ERK (1:1000, 4696, Cell Signaling Technology), MEK (1:1000, 4694, Cell Signaling Technology), Phospho-MEK (1:1000, 9154, Cell Signaling Technology), β-Actin (1:3000, 4970, Cell Signaling Technology), KRAS (1:1000, WH0003845M1, Sigma-Aldrich) and RAF1 (1:1000, 610152, BD Bioscience).

### Stable cell line generation

293T cells were seeded at 70% - 90% confluency in 6-cm dishes and transfected with the expression vector and packaging vectors (psPAX2 and pMD2.G) at a 3:2:1 ratio using Lipofectamine 2000 Transfection Reagent (Thermo Fisher Scientific). After 48 h, conditioned medium containing viruses was collected and filtered through 0.45-μm filters (Millipore). Target cells were seeded at 30% confluency in 6-well plates and infected with 500 μl of virus in the presence of 8 μg/ml Polybrene (Millipore) for 24 h. Inducible RAF dimer cell lines were generated by transducing cells with pLIX_403-BRAF-DmrC and either pInducer20-Blast-CRAF-DmrA for BRAF-CRAF heterodimer or pInducer20-Blast-BRAF-DmrA for BRAF homodimer. The cells were then selected with 2 μg/ml puromycin (Thermo Fisher Scientific) or 30 μg/ml blasticidin (Thermo Fisher Scientific) for 3 days.

### KRAS activation assay

Active KRAS (KRAS-GTP) was detected using the Active Ras Pull-Down and Detection Kit (Thermo Fisher Scientific). After cell lysis and protein quantification, 100 μg of cell lysates were incubated with 40 μg of glutathione S-transferase (GST)–RAF1 RAS-binding domain (RBD) and 100 μl of glutathione agarose resin in 1X lysis/binding/wash buffer for 1 h at 4 °C with gentle rotation, followed by three washes with 1X lysis/binding/wash buffer and elution with 80 μl of 2X SDS loading buffer. The samples were then analyzed by Immunoblotting using a KRAS specific antibody (Sigma-Aldrich).

### Cell viability and cell proliferation assay

Cells were seeded in 96-well plates at 2,000 cells per well in triplicates (at minimum) and treated with indicated concentrations of drugs for 72 h. For doxycycline inducible experiments, cells were pre-treated with indicated concentrations of dox overnight prior to drug treatment. To induce the RAF dimer, cells were treated with 100nM A/C ligand together with the inhibitors for 72 h. After treatment, the plate was equilibrated at room temperature for 30 min. Then, 100 μl of CellTiter-Glo Reagent was added to each well, and the plate was incubated on a shaker at room temperature for 10 minutes. Luminescence was measured using the GloMax plate luminometer (Promega).

### Cloning and plasmids

To generate doxycycline(dox) inducible constructs, genes were first cloned into the pENTR/D-TOPO vector using the pENT/D-TOPO Cloning Kit (Thermo Fisher Scientific). Gene inserts were then subcloned into either the pInducer20-Blast vector (a gift from Jean Cook, Addgene #109334) or the pLIX_403 vector (a gift from David Root, Addgene #41395) using the Gateway LR Clonase II enzyme mix (Thermo Fisher Scientific). For inducible RAF dimer constructs, DmrA or DmrC sequences were added to the C-terminus of RAF with a linker (5′-GGGGGAGGAGGATCT-3′). For the live cell luciferase experiments, KRAS mutants were cloned into the pBiT1.1-N (LgBiT) vector (Promega), while CYPA, BRAF, CRAF-RBD (amino acids 52–131), CRAF-RBDCRD (amino acids 52–188), BRAF-DmrA, BRAF-DmrC, CRAF-DmrA and CRAF-DmrC were cloned into the pBiT2.1-N (SmBiT) vector (Promega). For dox controlled expression of ligand induced RAF dimer vectors, BRAF-DmrC was cloned into the pLIX_403 vector, while BRAF-DmrA and CRAF-DmrA were cloned into the pInducer20-Blast vector. The DmrA and DmrC sequences are provided in the supplementary materials. To generate BRAF^3X^ and CRAF^3X^ mutants, L232, F237, and E275 in the linker-CRD of BRAF, as well as L136, F141, and K179 in the linker-CRD of CRAF, were mutated to alanine. Mutagenesis was performed using the QuikChange II XL Site-Directed Mutagenesis Kit (Agilent) following the manufacturer s instructions.

### Clonogenic assay

Cells were seeded in 12-well plates at 1 × 10^5^ cells per well and treated for 7 to 10 days. Media containing drugs was refreshed every 3 days. Cells were fixed with 1 ml of pre-chilled methanol for 10 min, then stained with 500 μl of 0.5% crystal violet solution for 30 min at room temperature. After staining, plates were thoroughly washed with water and scanned.

### Live cell split-luciferase assay

KRAS variants were cloned into the NanoBiT vector 1.1 (Promega) and CYPA or the RBD domain of CRAF (amino acids 52–131) were cloned into the NanoBiT vector 2.1 (Promega). 293T cells were seeded into white 96-well tissue culture plates (Corning) at 2 × 10^4^ cells per well in triplicates (at minimum). Cells at each well were transfected with equal amounts (50 ng) of LgBiT vectors and SmBiT vectors using Lipofectamine 2000 Transfection Reagent (Thermo Fisher Scientific) or JetOPTIMUS Transfection Reagent (Polyplus). For the measurement of RAS–CYPA or RAS-RBD complex formation as a function of inhibitor concentration, the cells were treated with daraxonrasib or DMSO for 2 h, NanoGlo luciferase substrate (Promega, N2011) was added, and the activity of reconstituted NanoBiT luciferase was detected in a GloMax plate luminometer (Promega). In the KRAS:RAF-dimer experiments, 50 ng of LgBiT-KRAS and 25 ng of each SmBiT-RAF-DmrA/DmrC vectors were co-transfected. RAF dimerization was induced by simultaneous addition of 100 nM A/C ligand at the time of transfection. To measure the KRAS:CYPA interaction in the presence of BRAF mutants, 5 ng of the pLIX_403-BRAF vectors were co-transfected with 50 ng of LgBiT-KRAS and 50 ng of SmBiT-CYPA. Cells were pre-treated with the indicated concentrations of dox overnight to induce BRAF mutant expression. To evaluate the KRAS:CYPA interaction in the context of RAF dimer, 5 ng of the pLIX_403-BRAF-DmrC and 5 ng of the pInducer20-CRAF-DmrA vectors were co-transfected with 50 ng of LgBiT-KRAS and 50 ng of SmBiT-CYPA. Cells were pre-treated with 10ng/ml dox and 100 nM A/C ligand overnight to induce the BRAF-CRAF dimer formation. 24 hours post-transfection, the culture medium was replaced with 100μl of Opti-MEM Medium (Thermo Fisher Scientific) containing the indicated concentrations of drugs for 2 h. Then, 25μl of Nano-Glo Live Cell Reagent (Promega) was added to each well, and the plates were gently mixed by hand. The bioluminescence signal was measured immediately using the GloMax plate luminometer (Promega).

For the measurement of RAS–CYPA binding as a function of time, cells were exchanged into cell culture medium containing Nano-Glo Endurazine live-cell substrate (Promega) and allowed to equilibrate at 37 °C for 1 h. Cells were then transferred to a SpectraMax iD5 plate reader (Molecular Devices) equilibrated at 37 °C. After measuring the baseline of the signal, inhibitor was added, and the luminescence was recorded continuously.

### Chemical ligand induced RAF dimerization

The inducible RAF-DmrA/C heterodimer system has been previously described and was obtained from Takara Bio. The DmrA or DmrC binding domains were fused to the C-terminus of RAF protein using a flexible linker (GGGGS). Thus, addition of the A/C ligand induces RAF-dimerization via the DmrA and DmrC domains (see schematic in Supplementary Fig. 5A). Two chemically inducible RAF dimerization systems were constructed to answer different experimental questions. To evaluate the effect on KRAS:RAF-dimer interaction, RAF-DmrA or RAF-DmrC were tagged with small-bit luciferase (at the N terminus), whereas KRAS was tagged with large-bit luciferase (for a schematic describing the split-luciferase system see Supplementary Fig. 1E, F). When KRAS and RAF form a complex the luciferase enzyme is reconstituted, enabling detection of the complex via bioluminescence. In turn, treatment with the A/C ligand enables the determination of the effect of ligand-induced dimerization on the ability of daraxonrasib to displace RAF dimers from mutant KRAS in live cells. To determine the effect of ligand induced RAF dimerization on the ability of daraxonrasib to recruit CYPA to KRAS, or the effect of RAF dimerization on the ability of daraxonrasib to inhibit proliferation, the RAF-DmrA/C chimeric proteins (without the luciferase bits) were expressed under a dox-inducible promoter in 293T cells or KRAS mutant cancer cell lines. Cells stably expressing these proteins were pre-treated with 100 nM A/C ligand (Takara Bio) overnight to induce RAF dimerization before drug treatment. Of note, in both the split-luciferase and the dox-inducible ligand-induced dimerization systems, partial RAF dimerization can occur even in the absence of A/C ligand (due to the presence of a KRAS mutation, which stimulates such dimers). This likely underestimates the effect of A/C ligand effects in our studies. In an orthogonal approach we relied on the CRD mutants to rescue the effect of A/C ligand induced dimerization.

## Acknowledgments

We are deeply grateful to the patients and their families for the generous donation of clinical specimens used in this study. We thank Neal Rosen and David Solit for their valuable insights on this study; and Megan Mroczkowski for discussing this work throughout its stages and for reviewing the manuscript.

## Author contributions

P.L. conceived and supervised the study. B.S., L.F.Y, and P.L. designed experiments and analyzed data. A.C-N., F.H., D.J.R., A.V., and A.W. performed cell-based and/or biochemical experiments. B.M.W., S.R.P., A.I.S., I. G-L, D.S.H., R.Y., and K.C.A. provided clinical specimens. S.K., E.A., K.K.L, A.H. and J.A.S. coordinated the sequencing and analysis of clinical specimens. A.D. provided experimental troubleshooting and key reagents. B.S., L.F.Y, and P.L. were the main writers of the manuscript. All authors reviewed and edited the manuscript as needed.

---

**Supplementary Fig 1.**
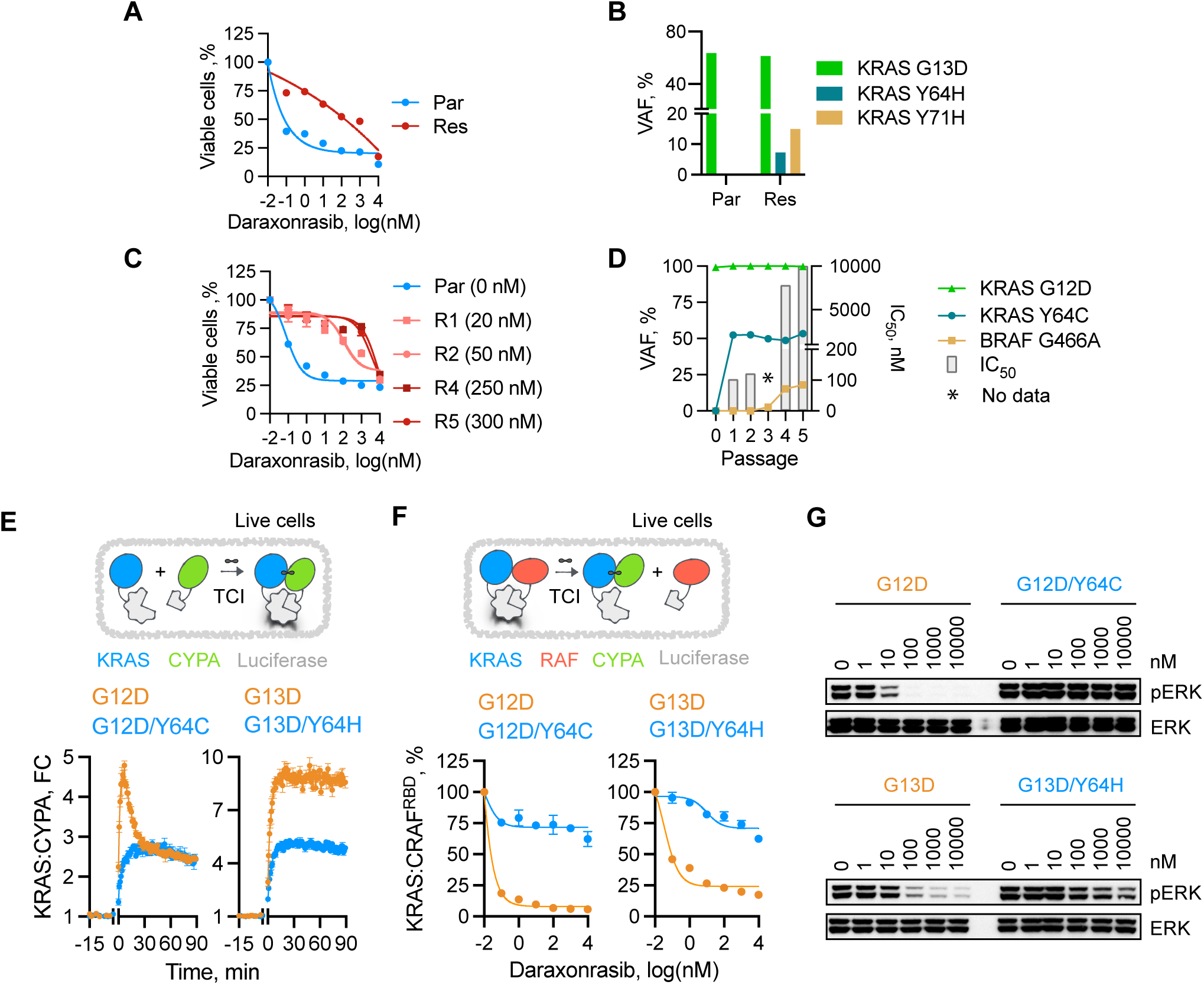
Identification and characterization of secondary KRAS mutations leading to daraxonrasib resistance. **A**) KRAS^G13D^ mutant cells (LoVo) were cultured longitudinally with increasing concentrations of daraxonrasib (or its pre-clinical analogue RMC-7977) to a final concentration of 100 nM. Parental (Par) and resistant (Res) cells were treated with daraxonrasib for 72 h to determine the effect on cell proliferation. **B**) Variant allele frequencies of KRAS mutations identified in the indicated cells. **C**) KRAS^G12D^ mutant cells (AsPC-1) were cultured longitudinally with increasing concentrations of daraxonrasib to a final concentration of 300 nM. Parental and serial resistant cell clones (R1-5) were treated with daraxonrasib for 72 h to determine the effect on cell proliferation. The daraxonrasib selection concentration is shown in parenthesis. **D**) Variant allele frequencies of the alterations identified in serial passages along with their respective daraxonrasib IC_50_ values. **E**) Live cells expressing a split luciferase reporter detecting the complex between CYPA and KRAS mutants were treated with 100 nM daraxonrasib and assayed over time, before and after treatment. FC: fold change. **F**) Live cells expressing a split luciferase reporter detecting the complex between the RBD of CRAF and KRAS mutants were treated with daraxonrasib for 2 h. **G**) Extracts from HEK293T cells expressing the indicated single- or Y64 double-mutants treated with daraxonrasib (2 h) were analyzed to determine the effect on ERK activation. In **A**, **C**, **E**, **F**, n = 3, mean ± s.e.m. A representative of two independent experiments is shown in **G**.

**Supplementary Fig 2.**
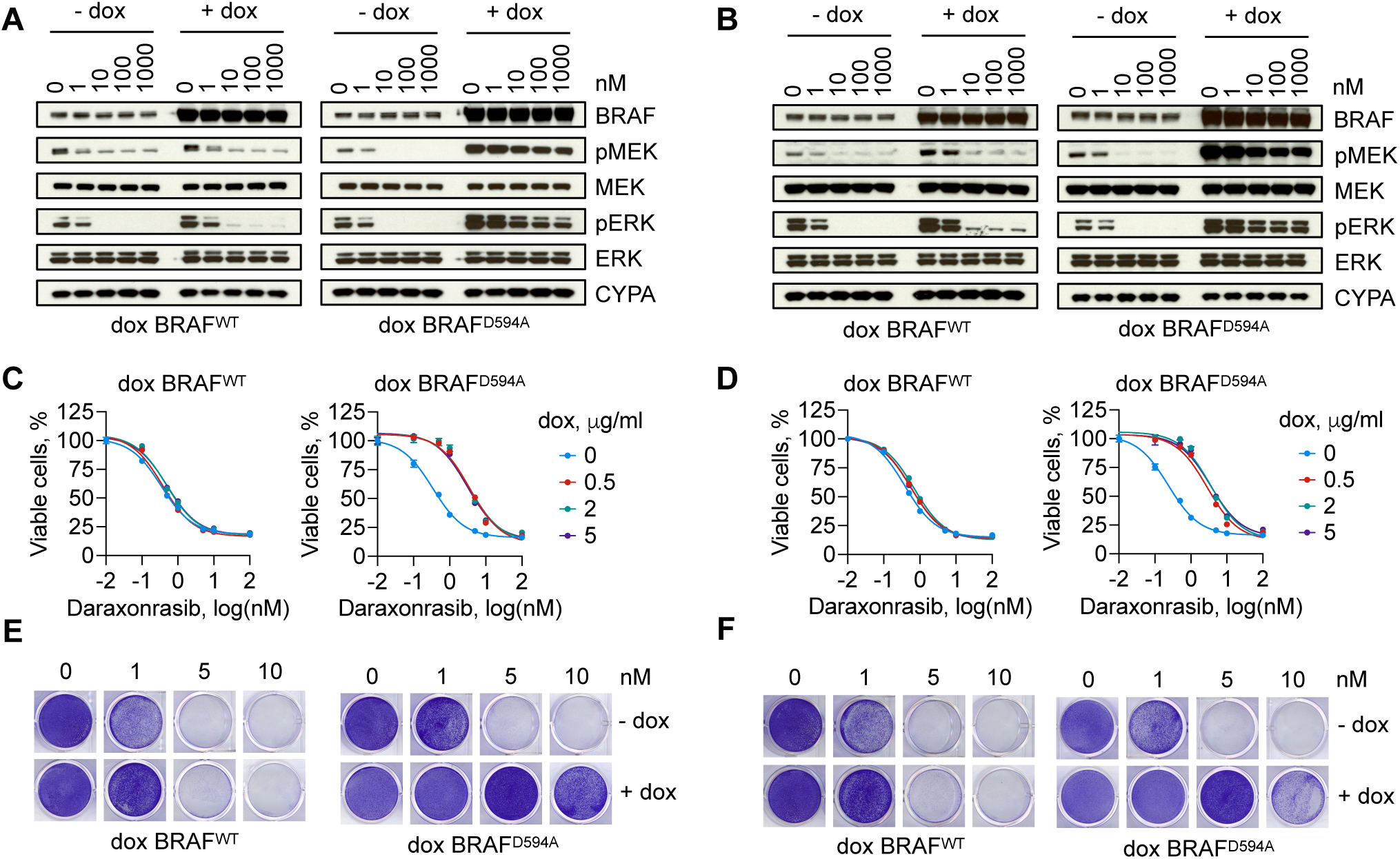
Kinase-dead BRAF antagonizes the effect of daraxonrasib treatment. **A**-**F**) KRAS^G12V^ mutant lung (H727, **A**, **C**, **E**) or colorectal (SW620, **B**, **D**, **F**) cancer cells expressing wild type or D594A mutant BRAF under a dox-inducible promoter were treated increasing concentrations of daraxonrasib for 2 h to determine its effect on ERK signaling (**A**, **B**), for 72 h to evaluate cell viability (**C**, **D**), or for 7 days to perform a clonogenic assay (**E**, **F**). A representative of two independent experiments is shown in **A**, **B**, **E**, **F**. In **C**, **D**, n = 3, mean ± s.e.m.

**Supplementary Fig 3.**
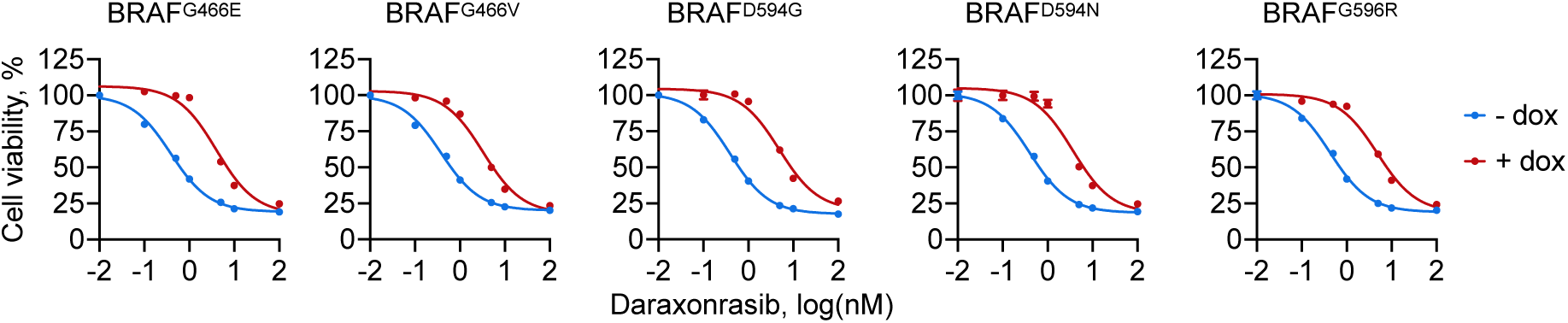
Class 3 BRAF mutants attenuate the antiproliferative effect of treatment. KRAS^G12V^ mutant (SW620) cells expressing the indicated hypoactive BRAF mutants under a dox-inducible promoter were treated with daraxonrasib for 72 h to determine the effect on cell viability. n = 3, mean ± s.e.m.

**Supplementary Fig 4.**
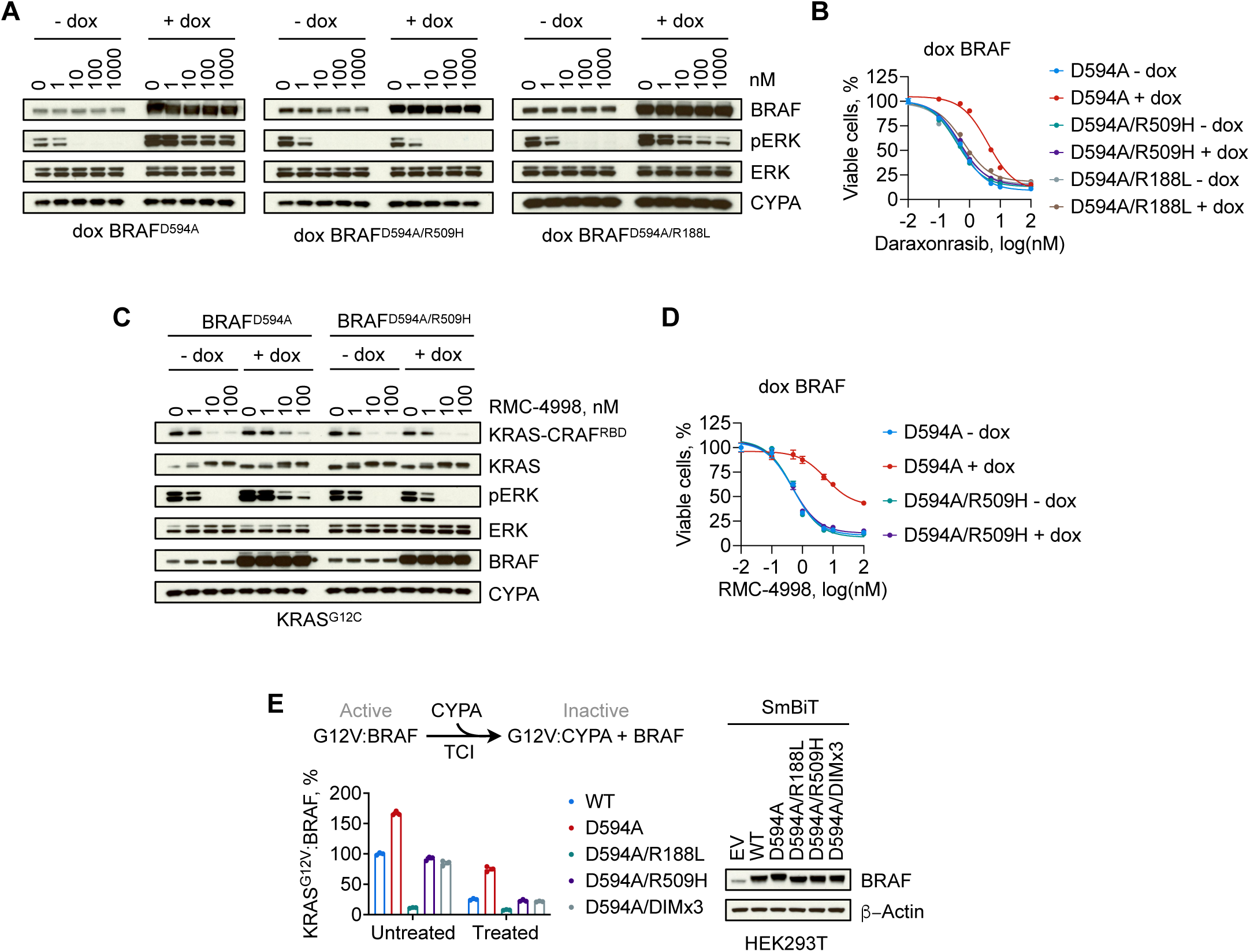
Kinase-dead BRAF drives resistance through RAS-dependent dimerization. **A**, **B**) KRAS^G12V^ mutant (SW620) cells expressing the indicated BRAF mutants under a dox-inducible promoter were treated with increasing concentrations of daraxonrasib for 2 h to determine the effect on ERK signaling (**A**) or for 72 h to determine the effect on cell viability (**B**). **C**, **D**) KRAS^G12C^ mutant (H358) cells expressing the indicated BRAF mutants under a dox-inducible promoter were treated with increasing concentrations of the G12C(ON) inhibitor RMC-4998 for 2 h to determine the effect on the interaction of KRAS with the RBD of CRAF (which is a proxy for KRAS-GTP levels) and on ERK phosphorylation (**C**), or for 72 h to determine the effect on cell viability (**D**). **E**) Live cell split-luciferase assay detecting the effect of daraxonrasib (100 nM, for 2 h) on the interaction between KRAS^G12V^ (tagged with LgBiT luciferase) and the indicated BRAF variants (tagged with SmBiT luciferase). DIMx3: R509H/L515G/M517W (triple dimer interface mutation). A representative of two independent experiments are shown in **A** and **C**. In **B**, **D**, **E**, n = 3, mean ± s.e.m.

**Supplementary Fig 5.**
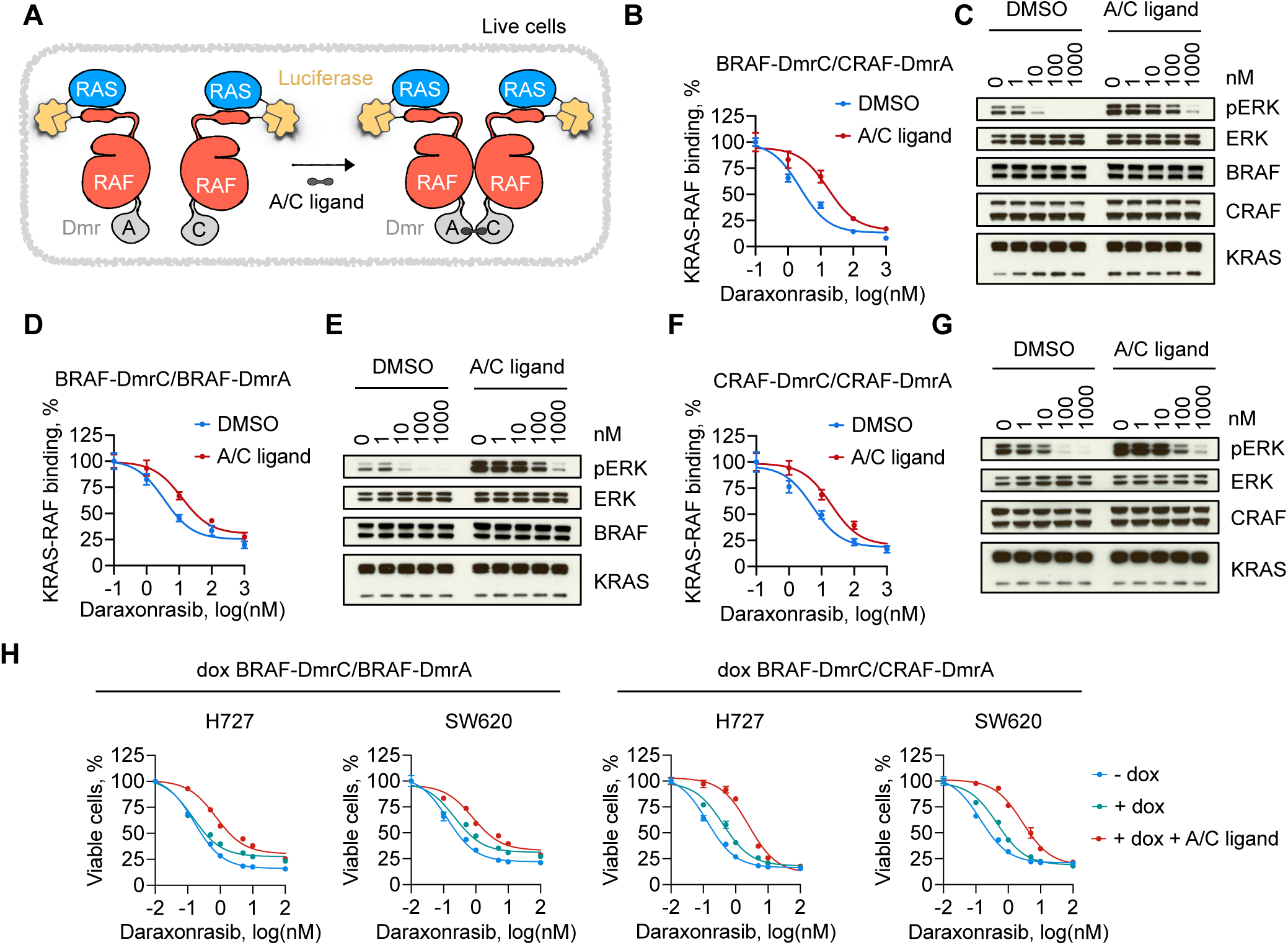
Ligand-induced RAF dimerization antagonizes the effect of treatment on target engagement, signaling and proliferation. **A**) A schematic of the chemical ligand-induced dimerization system. When determining the effect of dimerization on RAS-RAF interaction (**B**-**G**), these proteins were respectively tagged with large (LgBiT) or small (SmBiT) bit luciferase, as shown. When determining the effect on KRAS-CYPA interaction or cell proliferation (**h**), the RAF-Dmr chimeric proteins (without the split-luciferase) were stably expressed in KRAS mutant cancer cells under a dox-inducible promoter. **B**, **C**) HEK293T cells expressing LgBiT-KRAS^G12V^ and SmBiT-BRAF-DmrC/SmBiT-CRAF-DmrA chimeric proteins were treated with increasing concentrations of daraxonrasib (2 h) and either DMSO (control) or A/C ligand (to induce dimerization). The effect on KRAS-RAF interaction (**b**) and on ERK signaling (**C**) are shown. **D**-**G**) As in **B** and **C** but with the indicated SmBiT-RAF-DmrA/C chimeras. **H**) KRAS^G12V^ mutant lung (H727) or colorectal (SW620) cancer cells expressing the indicated chimeric constructs under a dox-inducible promoter were treated with daraxonrasib for 72 h to determine the effect on cell viability. A/C ligand (100 nM) was added as indicated. In **B**, **D**, **F**, **H**, n = 3, mean ± s.e.m. A representative of two independent experiments is shown in **C**, **E**, and **G**.

**Supplementary Fig 6.**
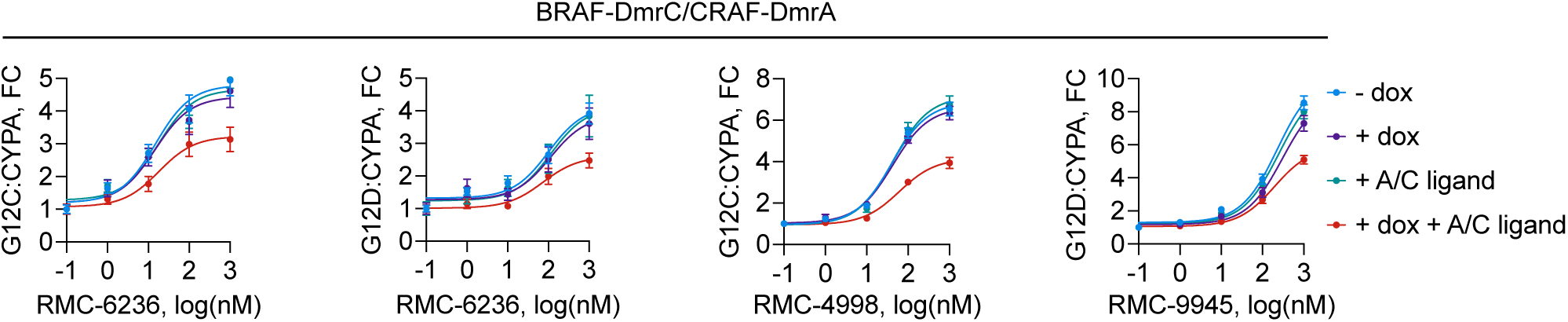
RAF dimerization attenuates the TCI-induced recruitment of CYPA to mutant KRAS. Live cell split-luciferase assay detecting the effect of dox-inducible expression of the indicated BRAF/CRAF ligand-induced dimerization constructs on the ability of the indicated TCIs to recruit CYPA onto mutant KRAS. FC: fold change. n = 3, mean ± s.e.m.

**Supplementary Fig 7.**
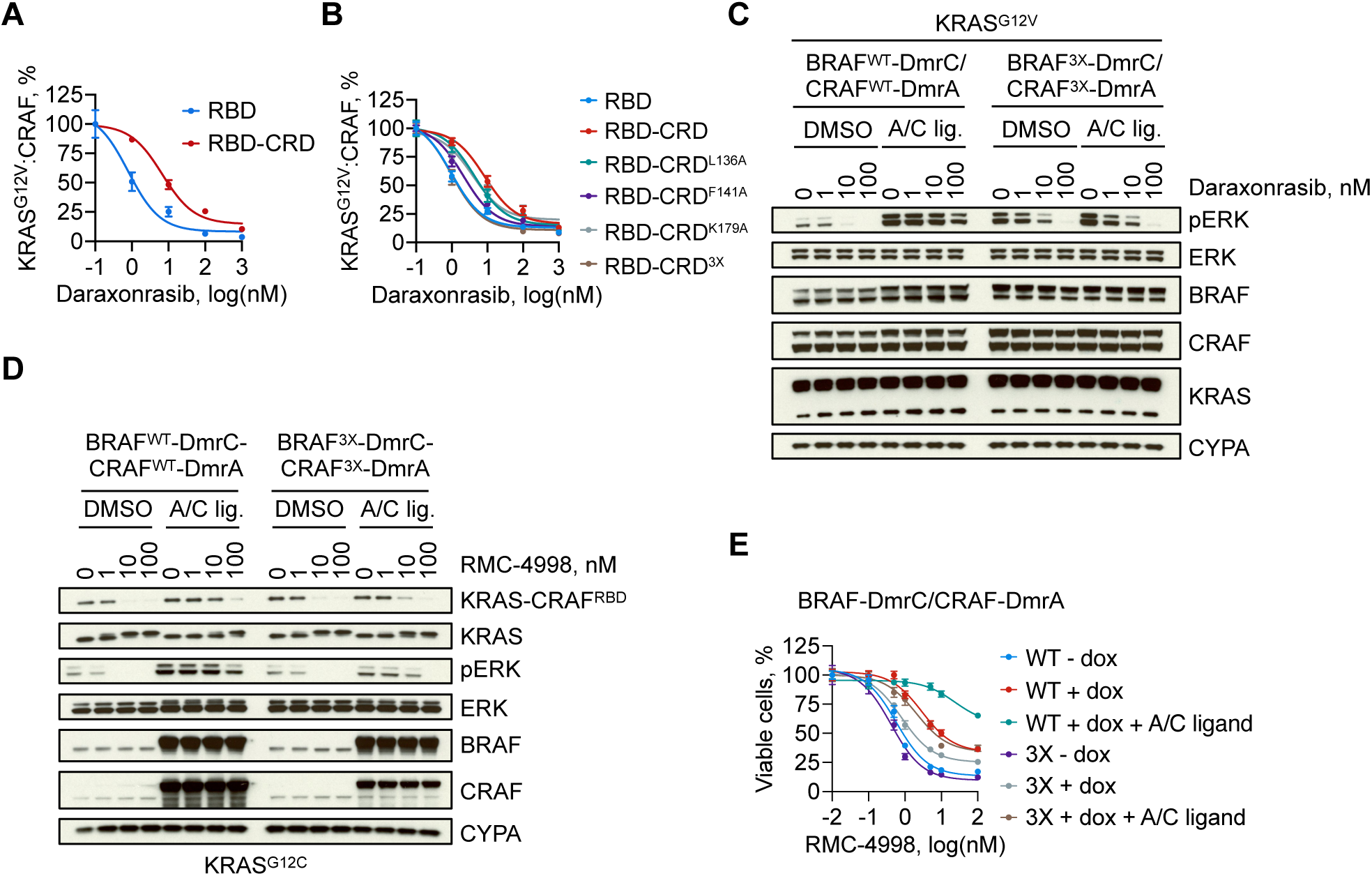
The cysteine-rich domain of RAF mediates the effect of dimerization on antagonizing daraxonrasib. **A**) Live cell split-luciferase assay detecting the effect of daraxonrasib treatment (2 h) on the interactions between mutant KRAS and the RBD or the RBD-CRD domains of CRAF. **B**) As in **A** but with the indicated mutants. RBD-CRD^3X^: L136A/F141A/K179A. **C**) HEK293T cells expressing the LgBiT-KRAS^G12V^ and SmBiT-BRAF-DmrC/SmBiT-CRAF-DmrA chimeric proteins were treated with increasing concentrations of daraxonrasib (2 h) and either DMSO (control) or A/C ligand (to induce dimerization). The effect on ERK phosphorylation is shown. **D**) KRAS^G12C^ mutant (H358) lung cancer cells, expressing the indicated ligand induced dimerization constructs under a dox-inducible promoter, were treated with the G12C(ON) TCI RMC-4998. Note how A/C ligand induced dimerization attenuates TCI binding to KRAS^G12C^ (KRAS electrophoretic motility shift), the inhibition of KRAS-CRAF^RBD^ interaction and the inhibition of ERK phosphorylation, as well as the reversal of these effects with the CRD mutant (3X). In **A**, **B**, **E**, n = 3, mean ± s.e.m. A representative of two independent experiments is shown in **C** and **D**.

